# Pvf1-PvR-mediated crosstalk between the trachea and the gut guides intestinal stem cell migration to promote gut regeneration

**DOI:** 10.1101/2024.09.02.609652

**Authors:** D. Mackay, A. John, C.F. Christensen, R. Loudhaief, A.B. Tanari, M. Rauzi, J. Colombani, D.S. Andersen

## Abstract

In adult tissues, stem cells (SCs) reside in specialized niches, where they are maintained in a quiescent state until activated by injury. Once activated, they migrate towards injured sites, where they proliferate and differentiate to replenish lost or damaged cells. Although effective tissue repair relies critically on the ability of SCs to reach and populate damaged sites, mechanisms guiding SCs towards these sites are not well understood. This is largely due to the technical challenges involved in monitoring SC dynamics in real time *in vivo*. Here, we devised an experimental framework that allows for real-time tracking of the spatiotemporal dynamics of intestinal SCs (ISCs) during the early phases of gut regeneration. Our data show that ISC migration is rapidly induced following injury and precedes ISC divisions and differentiation. We identify the Drosophila PDGF-VEGF-related receptor, Pvr, as a critical regulator of the migratory response to epithelial damage. ISC-specific Pvr depletion strongly suppresses ISC migration towards affected sites as well as the regenerative response. We further show that the Pvr ligand, PDGF-VEGF-related factor 1 (Pvf1), is produced by the trachea/vasculature in response to intestinal damage and acts as a guidance signal to direct ISC migration towards affected areas. Our work highlights a critical role of gut-trachea/vasculature crosstalk in guiding ISC migration during regeneration. As neovascularization of injured sites is a key feature of tissue repair in both flies and mammals, these findings could be relevant to regenerative processes in a wide range of adult tissues.

## Introduction

The inherent ability of adult stem cells (SCs) to assume a migratory behavior in response to appropriate stimuli has gained a lot of attention in recent years due to the potential of exploiting such behavior for wound repair and regenerative medicine. Through their capacity to self-renew and generate committed daughter cells, SCs produce a continuous source of differentiated cells essential for the maintenance and repair of adult tissues. While the regulations underpinning SC divisions and differentiation are increasingly well understood, the signals that guide SCs towards injured areas and ensure their correct positioning within the tissue remains relatively unexplored. Importantly, active migration of somatic SCs is thought to constitute an important component of the regenerate response in almost all adult tissues and has been particularly well studied in the context of skin and bone fracture healing processes (Dekoninck and Blanpain 2019; Gonzales and Fuchs 2017). Studies in mice coupling intravital imaging with SC labelling have shown that hair follicle SCs (HFSCs) and interfollicular SCs (IFESCs) are actively recruited from different regions of the skin to the wounded area and participate in epidermal repair (Aragona et al. 2017; Park et al. 2017). Likewise, efficient repair of bone fractures depends on the recruitment and expansion of perivascular mesenchymal skeletal stem and progenitor cells, which constitute a source of mature osteoblasts (Bohm et al. 2019; Su et al. 2018; Tang et al. 2009). To date, most signals reported to regulate cell migration were identified based on their capacity to stimulate or inhibit migration *in vitro* and by manipulating one component of the time (de Lucas, Perez, and Galvez 2018). However, SC migration is likely to be regulated by the coordinated action of mechanical cues and biochemical signals released from the SC niche and nearby organs. Furthermore, the migratory modes of SCs *in vivo* deviate substantially from those observed *in vitro* highlighting the importance of studying the migration of SCs in their native environment (Friedl et al. 2012; de Lucas, Perez, and Galvez 2018).

The Drosophila midgut provides a powerful model for studying adult SC-mediated regenerative processes. The fly intestine is made up of epithelial monolayers of enterocytes (ECs), which are aligned on their basal sides on an extracellular matrix referred to as the basement membrane (BM). The only active stem cells in the gut are the intestinal SCs (ISCs), which are embedded basally throughout the epithelium and divide non-symmetrically to give rise to daughter ISCs and transient progenitor cells, enteroblasts (EBs) or enteroendocrine progenitor cells (EEPs), that are destined to differentiate into ECs or EE cells (EECs), respectively (Biteau and Jasper 2014; Guo and Ohlstein 2015; Zeng and Hou 2015). To maintain intestinal homeostasis, a complex network of signals emanating from the ISC niche (ECs, EECs, EBs, and visceral muscles (VMs)) ensures that cell loss is tightly coupled with ISC-dependent proliferation and cell fate decisions (Biteau and Jasper 2011; Jiang et al. 2011; Liang et al. 2017; Lin, Xu, and Xi 2008). In response to intestinal infections, augmented ISC divisions and accelerated tissue turnover rates ensures that dying ECs are replaced. In addition to gut resident cells, the tracheal system, an oxygen-delivering network functionally equivalent to the mammalian blood vessels, constitute an important component of the regenerative response (Perochon et al. 2021; Tamamouna et al. 2021). The terminal tracheal cells (TTC; equivalent to the mammalian tip cells) infiltrate the adult gut and respond to reactive oxygen species (ROS) emanating from damaged cells by triggering increased tracheole branching, which is essential for driving ISC divisions and intestinal repair (Perochon et al. 2021; Tamamouna et al. 2021). Notably, a recent study showed that inhibiting actin dynamics in ISCs, and hence their capacity to migrate, impairs damage induced ISC divisions, suggesting that ISC migration and divisions are intimately linked processes (Hu et al. 2021). Here we use transient exposure to DSS or the highly pathogenic *Pseudomonas entomophila* (P.e) gram-negative bacteria to introduce local/restricted damage of the gut epithelium and live imaging of wholemount guts to track the migration of ISCs towards affected areas. Our data show that ISC migration is rapidly induced following exposure to DSS or P.e and results in the formation of ISC clusters at damaged areas of the gut epithelium. We further show that PDGF-VEGF-related factor 1 (Pvf1), produced by the trachea/vasculature in response to intestinal damage, acts as a guidance signal to stimulate ISC migration and gut regeneration.

In mammals, PDGF and VEGF both act as chemoattractants with essential roles in tissue repair. The PDGF-BB/PDGFR axis function as a potent stimulator of a variety of cells *in vitro*, and the direct application of PDGF-BB to wounds has been shown to recruit fibroblasts and improve healing (Bohm et al. 2019; He et al. 2022; Lin et al. 2014; Massberg et al. 2003; Mishima and Lotz 2008; Nedeau et al. 2008; Pierce et al. 1989; Pierce et al. 1988; Rajkumar et al. 2006; Wang et al. 2018). Nevertheless, the physiologically relevant source of PDGF-BB and its regulation during tissue repair is not known. In flies, the PDGFR-VEGFR-related receptor, PvR, is a critical regulator of developmentally controlled cell migrations (Cho et al. 2002; Duchek et al. 2001; Heino et al. 2001; Janssens, Sung, and Rorth 2010; McDonald, Pinheiro, and Montell 2003; Wood, Faria, and Jacinto 2006). Interestingly, PvR and its ligand, PDGF-VEGF-related factor 2 (Pvf2), are both expressed in ISCs, albeit the role of Pvf2-PvR signaling in intestinal regeneration remains controversial. Hence, one study showed that Pvf2 is required for the regenerative response triggered by administration of paraquat (Choi et al. 2008), while another found that Pvf2 and PvR are dispensable for the proliferative response triggered by intestinal infections (Bond and Foley 2012). Here we show that PvR, but not Pvf2, is required for ISC migration and divisions triggered by intestinal infections and localized damage. We find that intestinal damage triggers Pvf1 expression in the gut-associated trachea, and that Pvf1 derived from tracheal cells act as stimulates ISC migration during tissue repair. Knocking down Pvf1 in tracheal cells impairs the ISC-driven regenerative response, highlighting the coupling between ISC migration and divisions. As neovascularization is a hallmark of both tissue repair in all adult organs, these findings are likely applicable to repair processes in a broad range of adult tissues.

## Results

To shed light on the role of Pvr signaling in mediating regenerative growth, we knocked down Pvr in ISCs and their immediate progenitors, the EBs, using the ISC/EB specific driver, *esg*^*ts*^*>*, or in ISCs alone using *esg*^*ts*^*>* in combination with EB-specific expression of the Gal4 inhibitor Gal80 (*esg*^*ts*^*::Gal4; Su(H)::Gal80*; hereafter referred to as *ISC*^*ts*^*>*) and examined the effect on infection-induced ISC divisions and tissue turnover rates. Consistent with a role of Pvr in regulating ISC dynamics, ISC-specific knockdown of Pvr strongly suppressed the increase in ISC proliferation associated with both DSS treatment and enteric infections (Figure 1a-c). Moreover, knockdown of Pvr in ISCs and EBs (*esg*>) strongly suppressed the infection-induced increase in proliferation and tissue turnover rates (Figures 1d-e). By contrast, Pvf2 depletion did not affect the accelerated tissue turnover associated with enteric infections (Figure 1e). Consistent with a role of Pvr in mediating regenerative growth, knockdown of Pvr in ISCs using two different RNAis increased the sensitivity to oral infection with the highly pathogenic *Pseudo entomophila* (Pe) bacterium (Figure 1f). As previously mentioned, ISC migration and divisions are thought to be closely interlinked processes. To analyze how ISC-specific Pvr knockdown affects actin dynamics in a regenerative context, we expressed RFP-tagged LifeAct in ISCs. As previously reported (Hu et al. 2021), global damage triggered the formation of highly dynamic actin-based lamellipodia within two hours of an oral infection (Figure 1g-h, movies 1-3). Strikingly, ISC-specific Pvr knockdown completely abolished the infection-induced formation of actin-based protrusion quantified as an increase in circularity (Figure 1g-k). Altogether, these data support an ISC-specific role of Pvr in promoting damage-induced actin-based protrusions previously reported to be essential for ISC migration.

**Figure 1:**
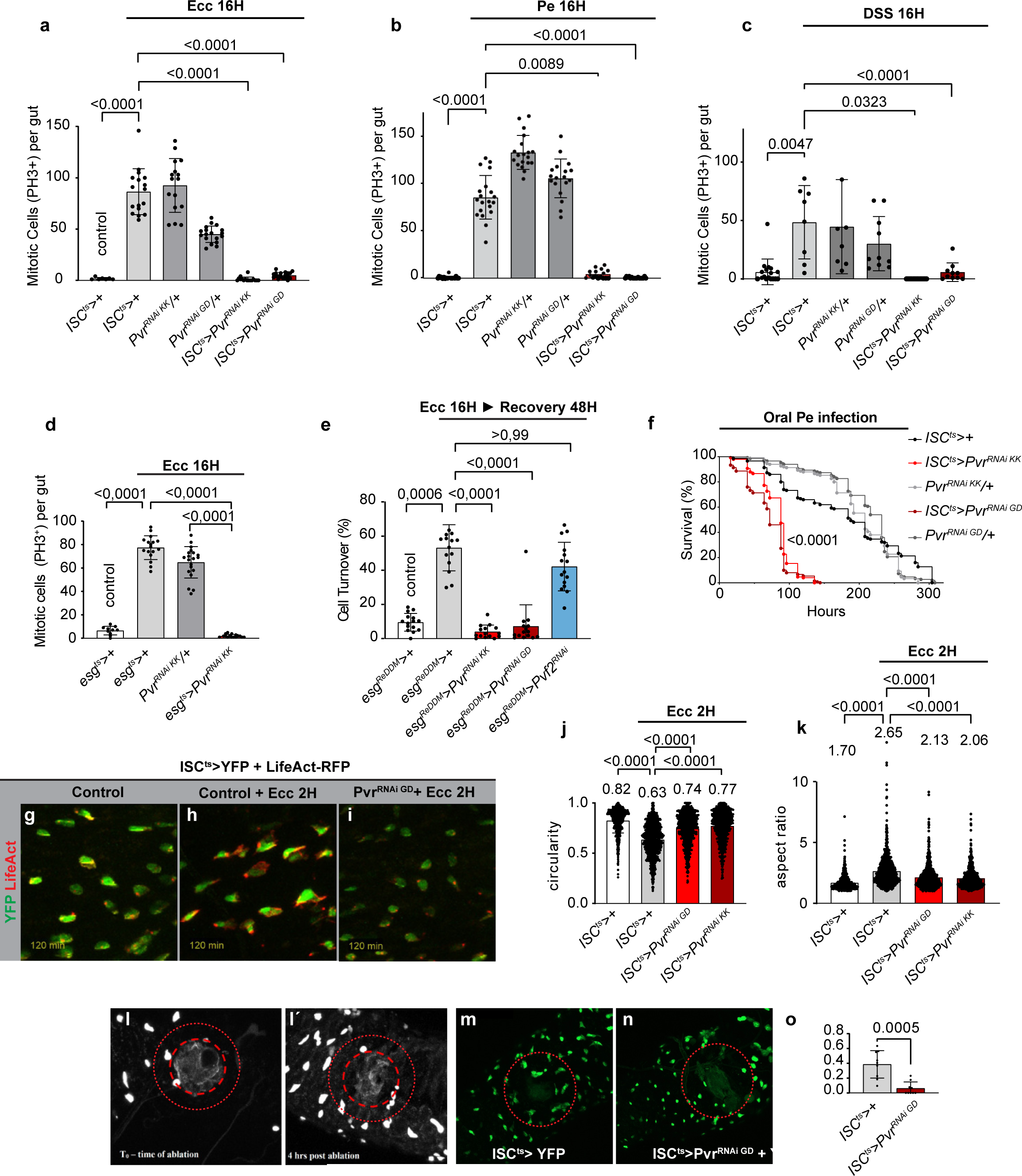
ISC-specific Pvr knockdown impairs the regenerative response and resistance to oral infection. (a-c) Quantification of PH3+ cells in midguts dissected from control flies or flies with ISC - specific RNAi-mediated Pvr knockdown (ISC^ts^>Pvr RNAi) subjected to oral infection with Ecc15 or Pe (a-b) or DSS treatment (c). (d) Quantification of PH3+ cells in midguts dissected from control flies or flies with ISC/EB-specific Pvr knockdown *esg*^*ts*^*>Pvr* RNAi subjected to oral infection with Ecc15. (e) ISC/EB-specific knockdown of Pvr, but not Pvf2, strongly suppresses infected-induced acceleration in tissue turnover rates. Membrane-tethered CD8::GFP and nuclear localized H2B::RFP is specifically co-expressed in stem and progenitor cells with the *esg>* driver. As progenitors differentiate into mature cells, loss of esg-Gal4 expression terminates further production of GFP and RFP. Differential stability of the fluorophores results in rapid degradation of GFP whereas RFP remains in differentiated cells for an extended duration of time. For all ReDDM experiments, RFP-positive/GFP-negative polyploid cells were scored as newly produced ECs. (f) Mated female flies with ISC-specific knockdown of Pvr display reduced survival to oral Pe infection. (g-j) To monitor actin dynamics in ISCs after oral infection with *Ecc15*, ISC^ts^> was used to drive YFP (to label ISCs in green) and LifeAct (to label F-actin in red alone (without (g) or with (h) infection, controls) or in combination with Pvr RNAi (i). Representative confocal images of dissected posterior midguts from control flies (**g-h**) and flies expressing *Pvr*^RNAi^ (**i**), showing that ISC-specific Pvr depletion disrupts the formation of actin-based protrusions triggered by oral infection, quantified as an increase in circularity (j) or a decrease in aspect ratio (k). (l-l’) High-powered lasers with two-photon microscopy were utilized to generate a localized ∼50 μm wound (red stippled circle). Confocal images of the R4 region dissected from control flies *at the time of ablation* (T_0_, l) and 4 hours later (T_4_, l’) with ISCs labelled in white. The number of migrated ISCs was quantified as the number ISCs within a radius of 100 μm of the wound (red dotted line) 4 hours after ablation (T_4_) subtracted by the number of ISCs already present at the time of ablation (T_0_). (m-o) Representative confocal images of guts dissected from control flies (**m**) and flies with ISC-specific Pvr knockdown *(n)* 4 hours after ablation with ISCs labelled in green with the number of migrated ISCs quantified in o. Significance was tested with a two-tailed Mann-Whitney test (n), or Kruskal Wallis with post-hoc multiple comparison analysis. Data are presented as mean values ±SD.

To obtain a condition for studying the directed movement of ISCs towards a localized area of damage, we next used pulsed two-photon infrared lasers to regenerate a localized wound of about 50 μm and live imaging of wholemount guts to monitor ISC migration. As previously reported, the toxicity associated with live imaging causes guts to deteriorate within a few hours of wounding, and hence, was not compatible with monitoring ISC migration in real time. Instead, we defined a radius of 100 μm around the ablation site and counted the number of ISCs at the time of ablation and 4 hours later. We observed an accumulation of ISCs 4 hours post-ablation around the wound site. According to previous reports and our own observations, ISCs do not divide within the first 4 hours of epithelial wounding (Hu et al. 2021). ISC-specific Pvr knockdown prevented the accumulation of ISCs at the wound, consistent with the role of Pvr in promoting ISC migration in response to epithelial damage (Figure 1l-o).

To device an alternative strategy allowing ISC migration to be followed in real time, we introduced a GFP-tagged E-Cadherin (E-Cad-GFP) in the background and used perturbation of membrane-associated E-Cad staining as a proxy for areas of epithelial damage. Following two hours of exposure to the damaging agent, DSS, we observed local/restricted regions of epithelial damage (see yellow star in Figure 2a-a’’’, movie 4). Intriguingly, while ISCs remained quiescent, in undamaged regions with regular membrane-associated E-Cad staining (Figure 2b-b’’’, movie 5), we observed directed movements of ISCs towards areas with disrupted E-Cad “pattern” resulting in the formation of ISC clusters within the time frame of 2-6 hours (Figure 2a-a’’’, movie 4). Similarly, we found that transient infection with the pore-forming Pe bacteria for two hours resulted in localized damage (visualized as loss of E-Cad) and the directed ISC movement of ISCs towards these areas within 2-4 hours of the infection (Figure 2c-d’’). Using these two strategies to induce local/restricted damage, we show for the first time that local wounding of the gut epithelium triggers a rapid response, where ISCs migrate over distances of several cell perimeters towards damaged areas forming ISC clusters (Figure 2c-d’’,g, movies 1 and 6). Importantly, we did not observe ISC divisions within this time frame. Consistent with a requirement for Pvr signaling in this process, we found that Pvr-depleted ISCs remained static and did not migrate or form ISC clusters in response to epithelial damage induced by Pe infection (Figure 2e-g, movies 7 and 8). To analyze the order of events involved in tissue regeneration, we used TransTimer, a genetic tool that allows ISC-specific expression of a short-lived GFP (half-life ∼ 2 hours) and a long-lived RFP (half-life ∼ 18,5 hours) to follow the maturation of ISCs (green and red) into EBs (red only, Figure 2h) (He et al. 2019). Notably, ISC-to-EB differentiation primarily occurred more than 10 hours after epithelial damage (Figure 2i-i’’, movie 9), indicating that ISC migration and the formation of ISC clusters precedes ISC division and maturation during regenerative growth.

**Figure 2:**
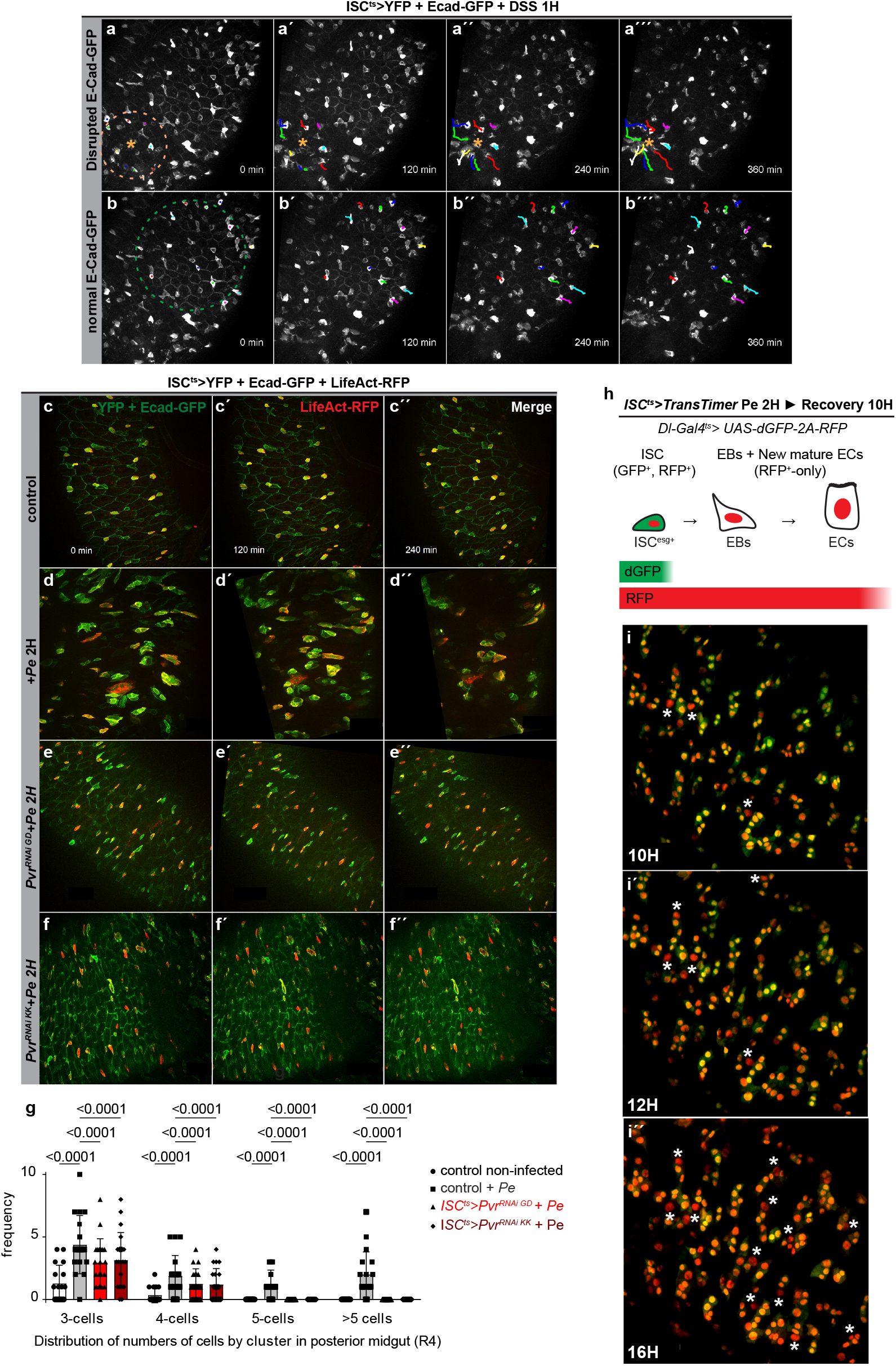
ISC-specific Pvr knockdown suppresses the migratory response to epithelial damage. (a-b’’’) Guts expressing E-Cad::GFP under its endogenous promoter to label cell-cell adhesion junctions (white, membrane-associated) with ISCs labelled in white (cytoplasmic white, ISC^ts^>YFP) dissected from flies that were fed DSS for two hours. Perturbation of membrane-associated E-Cad staining as a proxy for areas of epithelial damage and the migration of ISCs were traced towards damaged areas in real time using live imaging. (a-a’’’) ISCs show directed movements towards an area with disrupted E-Cad “pattern” (yellow star) over a period of 6 hours. (b-b’’’) In areas with intact E-Cad membrane association, ISCs remain relatively static and do not display directed movements. (c-g) Guts with ISC-specific expression of YFP to label ISCs (in green) and LiveAct-RFP (in red) to monitor actin dynamics dissected from uninfected flies (g, control) or flies exposed to Pe for two hours (h-i). Representative time points from real time recordings showing that ISC-specific Pvr knockdown suppresses ISC migration and the formation of ISC clusters, as quantified in (g) by counting the number of cells per cluster. The acquired movies were aligned in Fiji using “rigid body transformation to reduced movement triggered by muscle contractions. **(h)** Schematic diagram of the *TransTimer* system. dGFP and RFP is specifically co-expressed in stem cells with the *Dl*^*ts*^> driver. As progenitors differentiate into EBs, loss of *ISC* expression terminates further production of dGFP and RFP. Differential stability of the fluorophores results in rapid degradation of dGFP, which has a half-life of 2 hours, whereas RFP remains in differentiated cells for about 20 hours. (i-i’’) The TransTimer system was used to follow the differentiation of ISCs into EBs from 10 to 16 hours after infection. At 10 hours only few ISCs (red and green) have differentiated into EBs (red only, labelled by while stars), while 6 hours later many more ISCs have either differentiated into EBs or are in the process of transitioning (more red than yellow). Significance was tested with Kruskal Wallis with post-hoc multiple comparison analysis. Data are presented as mean values ±SD.

To decipher how Pvr signaling regulates actin dynamics, we ectopically expressed an activated form of Pvr, Pvr^CA^, and RFP-LifeAct in ISCs and monitored the effects on cell shape and actin-based protrusions. ISC-specific Pvr activation resulted in large-flattened ISCs that formed actin-based protrusions in a non-coordinated fashion, suggesting that Pvr signaling is sufficient to trigger lamellipodia formation (Figure 3a-c). Consistent with a critical role of Pvr in ISC differentiation (Figure 1e), ISC-specific Pvr activation is sufficient to accelerate tissue turnover rates in the absence of an infection (Figure 3d-e’’). Altogether these results show that ectopic Pvr signaling is sufficient to activate ISCs and promote their maturation into EBs.

**Figure 3:**
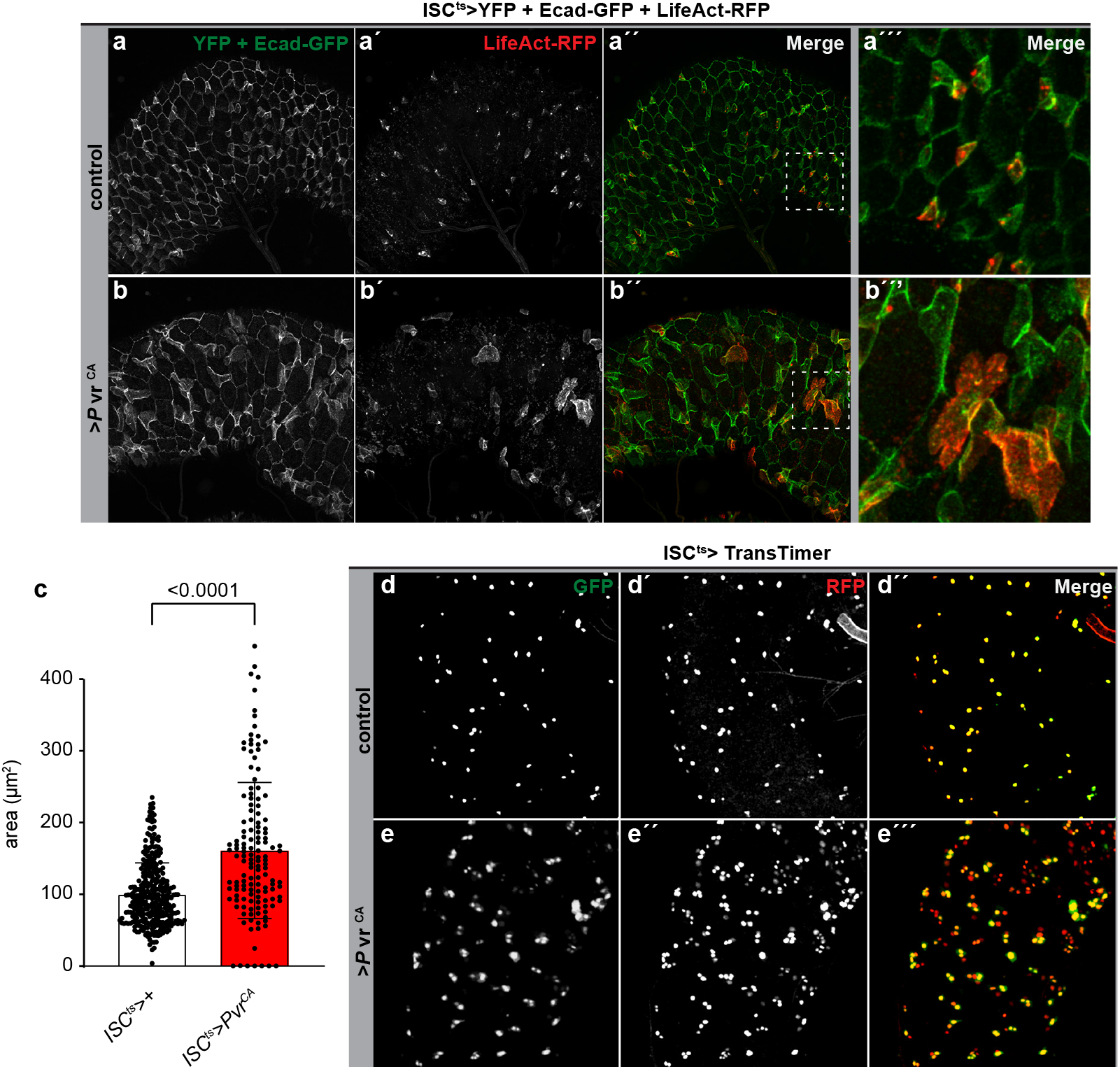
Ectopic Pvr activation is sufficient to activate ISCs and accelerate tissue turnover rates. (a-c) Representative confocal images of dissected guts expressing E-Cad::GFP under its endogenous promoter to label cell-cell adhesion junctions (in green) and ISC^ts^-driven expression of LifeAct (to label F-actin in red) alone (a-a’’’, control) or together with ISC-specific Pvr activation (b-b’’’, ISC^ts^>Pvr^CA^). Ectopic Pvr activation results in large-flattened ISCs that form actin-based protrusions in a non-coordinated fashion, quantified as an increase in cell size in (c). (d-e) The TransTimer system was used to follow the differentiation of ISCs into EBs in control guts (d-d’’) and guts with ISC-specific Pvr activation (e-e’’). Representative confocal images of dissected posterior midguts from control flies (**d-d’’**) and flies expressing *Pvr*^act^ (**e-e’’**) in stem cells using *ISC*^TransTimer^ tracing for two days. Pvr activation results in accelerated tissue turnover visualized by an increase in the number of EBs and ECs (red only cells, e-e’’) compared with control guts (d-d’’). Significance was tested with a two-tailed Mann-Whitney test. Data are presented as mean values ±SD.

Pvr signaling is activated upon binding of one of its three ligands, Pvf1-3. According to single-cell RNAseq data of the adult gut available at the FlyCellAtlas (https://scope.aertslab.org/#/FlyCellAtlas) (Li et al. 2022), Pvf3 is poorly expressed in the gut, while Pvf1 and Pvf2 show distinct expression patterns with Pvf2 being highly expressed in ISCs/EBs and Pvf1 in mature ECs. Consistent with this, a previous study found that Pvf2 expression is restricted to ISCs and EBs in the adult gut (Choi et al. 2008). As knockdown of Pvf2 in ISCs/EBs did not suppress the infection-induced acceleration in tissue turnover rates (Figure 1e), we focused our attention on Pvf1. Notably, Pvf1 was previously reported to act as a guidance signal for Pvr-mediated border cell migration (Duchek et al. 2001; Fulga and Rorth 2002; Janssens, Sung, and Rorth 2010; Zhou et al. 2022; Bianco et al. 2007). Consistent with a role of Pvf1 in promoting regenerative growth, Pvf1 expression (Figure 4e-f) and production (Figure 4a-d’’) was induced in several conditions associated with epithelial damage including oral infections with P.e (Figure 4a-b’’, d-d’’) or following ingestion of DSS (Figure 4c-c’’). While 16 hours of oral infection resulted in widespread induction of Pvf1-HA in VMs, ECs, and gut-associated trachea (Figure 4a-b’’), a shorter exposure (6 hours) to DSS or Pe induced Pvf1 expression and production in the gut-associated trachea and a subset of ECs that likely correspond to damaged areas of the epithelium (Figure 4c-d’’). To identify the source of Pvf1, we knocked it down in ECs, VMs and trachea. While knockdown of Pvf1 in ECs or VMs had no effect on proliferative response to oral infections (Figure S1a-b), trachea-specific Pvf1 depletion significantly reduced Ecc15 and Pe induced ISC divisions (Figure 4g-h) similar to what was obtained by knockdown of Pvf1 in ECs, VMs, and trachea altogether (Figure 4i), suggesting that trachea-derived Pvf1 might guide ISC migration during regeneration. Consistent with this, we found that trachea-specific knockdown of Pvf1 slowed down the migratory response to oral infection (Figure 4l-k), although ISC clustering was observed at later stages (data not shown). We noticed that trachea-specific knockdown of Pvf1 triggered a morphological change in ISCs that resulted in large-flattened cells that formed cellular protrusions in a non-coordinated fashion (Figure 4m-m’’’’, movies 10 and 11), reminiscent of that observed upon ISC-specific Pvr activation (Figure 3b-b’’’). Altogether, our results suggest that trachea-derived Pvf1 provides ISCs with a directional cue that is required for efficient migration of ISCs towards damaged areas. Two recent studies showed that the tracheal remodeling associated with damage of the gut epithelium is required to support the associated proliferative response (Perochon et al. 2021; Tamamouna et al. 2021). Importantly, knockdown of Pvf1 or Pvr in the trachea did not affect the infection induced increase in tracheal coverage (Figure 4j). On the other hand, ectopic expression of Pvf1 in tracheal cells was not sufficient to trigger surplus divisions in homeostatic conditions (data not shown), suggesting that the tracheal remodeling or additional damage induced signals is required to stimulate ISC migration and divisions. Altogether, our results show that damage of the gut epithelium triggers the expression of Pvf1 in the gut-associated trachea, which then acts as a chemoattractant to guide the migration of ISCs towards damaged areas.

**Figure 4:**
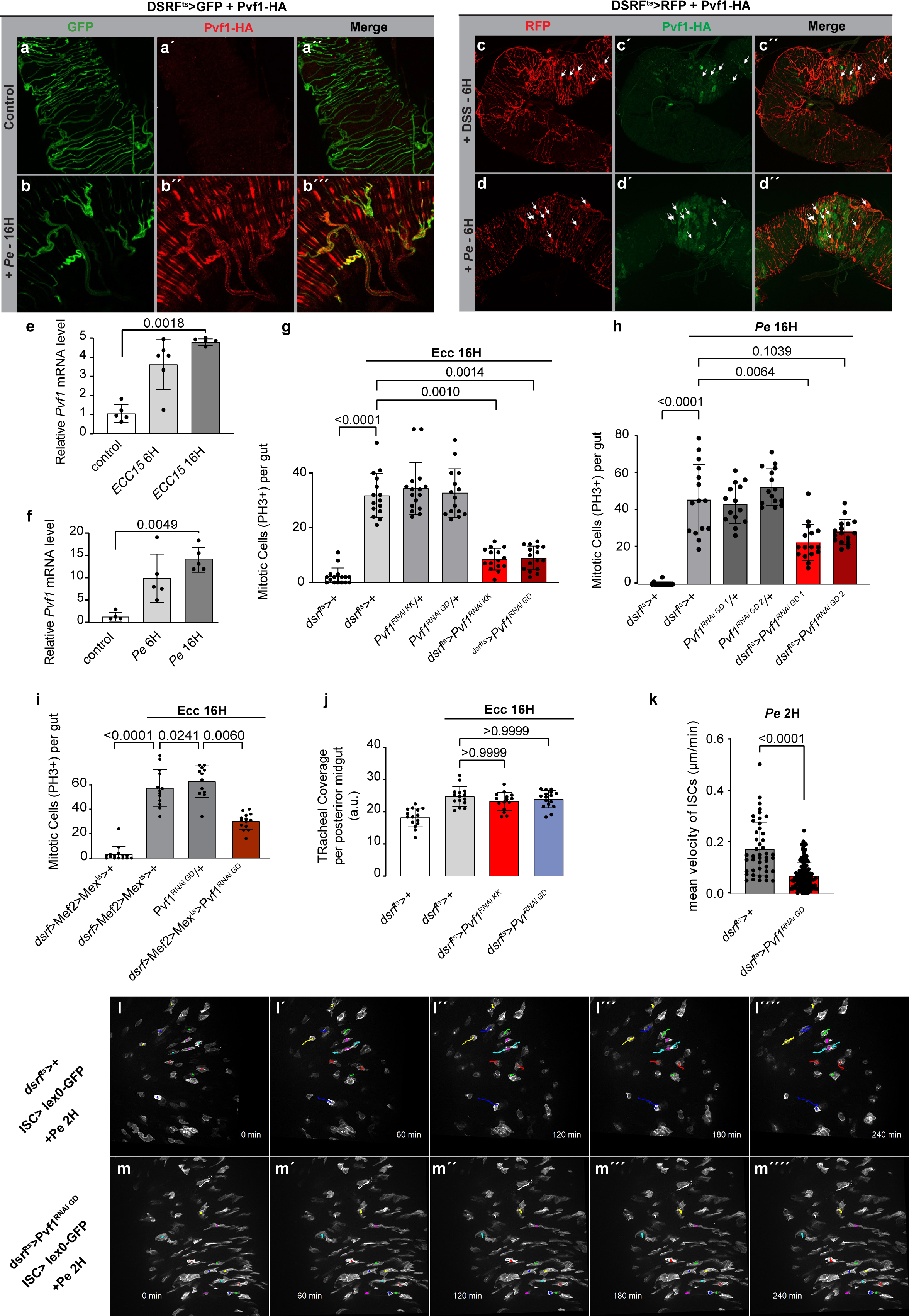
Pvf1 is required in the trachea to guide ISC migration and for an efficient regenerative response. (a-b’’) Representative confocal images of dissected posterior midguts from mock-treated flies or flies infected with Pe for 16 hours expressing *Pvf1*-HA under the control of its endogenous promoter, showing a widespread induction of Pvf1-HA in the gut epithelium, VMs, and trachea (**b’-b’’**). (c-d’’) Representative confocal images of dissected posterior midguts from mock-treated flies or flies with DSS (c-c’’) or infected with Pe (d-d’’) for 6 hours, shows that *Pvf1*-HA is mostly induced in a subset of ECs and tracheal cells at this stage (white arrows). RT-qPCR analysis on dissected control midguts showing that Pvf1 expression is induced at 6 and 16 hours after oral Ecc15 (e) and Pe (f) infection. (g-h) Quantification of PH3+ cells in midguts dissected from control flies or flies with trachea-specific RNAi-mediated Pvf1 knockdown subjected to oral infection with Ecc15 (g) or Pe (h). (i) Quantification of PH3+ cells in midguts dissected from control flies or flies with RNAi-mediated knockdown of Pvf1 in trachea, VM, and ECs subjected to oral infection with Ecc15 for 16 hours. (j) Quantification of tracheal coverage in guts dissected from control flies or Ecc15 infected flies with or without trachea-specific Pvf1 (red) and Pvr (blue) knockdown. Knockdown of Pvf1 or Pvr does not affect the infection-induced increase in tracheal coverage (visualized by *dsrf>* driven GFP). (l-k) Guts expressing GFP under the control of ISC^ts^>lexA-LexAop system to label ISCs (in white) were dissected from control flies (l-l’’’’) flies or flies with trachea-specific Pvf1 knockdown (*dsrf>*Pvf1 RNAi; m-m’’’’) after exposure to Pe for two hours. Representative time points from real-time recordings show that trachea-specific Pvf1 knockdown suppresses ISC migration (l-m’’’’), quantified as reduced velocity in (k). Significance was tested with a two-tailed Mann-Whitney test (k), or Kruskal Wallis with post-hoc multiple comparison analysis. Data are presented as mean values ±SD.

## Discussion

Identification of the signals the guide ISC migration during tissue repair has been hampered by the difficulty of following this process in real time *in vivo*. Here we provide the first real time recordings of ISC migration showing directed movements towards damaged areas of the gut epithelium. Taking advantage of the restricted/localized damage caused by brief exposure to the highly pathogenic Pe bacterium or damaging agents, we employed live imaging on cultured gut explants to monitor spatiotemporal ISC dynamics in the early phase of tissue regeneration. Our data show that ISC migration takes place within 2-6 hours of damaging the gut epithelium and precedes ISC divisions and differentiation. ISC-to-EB differentiation is observed 8-10 hours after epithelial damage when ISCs have already clustered. We further show that Pvr is a critical regulator of the migratory response associated with tissue regeneration, as ISC-specific Pvr depletion inhibits the formation of actin-based cellular extrusions and ISC migration in this condition. While Pvr knockdown impairs ISC migration, it also suppresses regenerative ISC divisions and maturation, highlighting the intimate link between ISC migration and divisions. Consistent with a role of Pvr in regulating actin dynamics, ectopic Pvr activation is sufficient to induce actin-based protrusions and a flattened morphology of ISCs in the absence of epithelial damage. Interestingly, Pvr activation was recently reported to trigger spreading and a similar lamellar morphology of hemocytes adhering to the surface of epidermal wounds in the Drosophila larvae (Tsai et al. 2022). One of the best characterized developmental functions of Pvr, is its role in collective border cell migration (Duchek et al. 2001; Fulga and Rorth 2002; Janssens, Sung, and Rorth 2010; Zhou et al. 2022; Bianco et al. 2007). Pvf1 released from the oocyte acts as a guidance signal to activate Pvr signaling in the leading cells, and ectopic Pvf1 expression is sufficient to redirect the migration of border cells (McDonald et al 2003). Pvr activation generates protrusions at the leading edge through the activation of Rac1 and its downstream effectors, Scar/Wave and the Arp2/3 complex (Zhou et al 2022). Consistent with a similar pathway operating in ISCs, we find that Pvr is required to activate lamellipodia formation and ISC migration in response to enteric infections. Our data also aligns with previous observations showing that ISC-specific knockdown of Rac1 or Arp3 suppresses lamellipodia formation triggered by enteric infections (Hu et al. 2021).

Crosstalk between trachea and gut is essential for mediating tissue regeneration. Two recent studies showed that damage of the gut epithelium triggers ROS-dependent remodeling of the trachea, which in turn is required to promote regenerative growth (Perochon et al. 2021; Tamamouna et al. 2021). Hence, it is tempting to speculate that ROS emanating from damaged epithelial cells induces Pvr1 expression in the gut-associated trachea, which subsequently guides ISCs towards injured sites. Interestingly, while Pvf1 is required in the trachea to promote regenerative growth, ectopic expression of Pvf1 in the trachea is not sufficient to trigger ISC divisions, suggesting that remodeling of the TTC might be necessary to deliver Pvf1 at damaged areas of the gut.

The same stimuli that trigger SC migration during tissue healing are often subverted into promoting migration and dissemination of cancer cells. Not surprisingly, abnormal PDGF and VEGF activities have been reported in a wide range of human tumors, and expression levels of PDGFRs/PDGFs correlate with invasiveness, tumors size, chemoresistance and poor clinical outcome (Pandey et al. 2023). Furthermore, studies on cancer cell lines have shown that PDGF-PDGFR signaling contributes to cancer cell dissemination *in vitro*. However, as the native microenvironment differs considerably from that of *in vitro* cell culture systems, it is essential to address the contribution of the PDGF/PDGFR axis to the dissemination of cancer cells *in vivo*. Here we show that ectopic Pvr activation triggers a partial Epithelial-mesenchymal transition (EMT) characterized by internalization and accumulation of the cell-cell adhesion proteins in the cytoplasm, altered F-actin dynamics and the formation of cellular protrusions accompanied by changes in cell morphology. The observation that Yorkie transformed ISC tumors express high levels of Pvf1 (Song et al. 2019), suggest that tumor-derived Pvf1 might potentiate cancer stem cell dissemination.

The acquisition of mesenchymal traits by SCs, essential for their mobility and for tissue regeneration, has been observed in a number of adult tissues, including the respiratory epithelium, a tissue that in structure and composition is similar to the fly gut (Forte et al. 2017; Haller et al. 2017). Hence, it is possible that crosstalk between the respiratory epithelium and the vasculature plays an identical role in guiding SC migration towards affected areas in this tissue. The partial EMT of ISCs during tissue regeneration must be complemented and balanced by the reverse process, Mesenchymal-Epithelial transition (MET), upon return to homeostasis. Understanding the mechanisms underpinning MET and the resolution of regenerative ISC clusters upon return to homeostasis will be an exciting future area of research.

---

**Figure S1:**
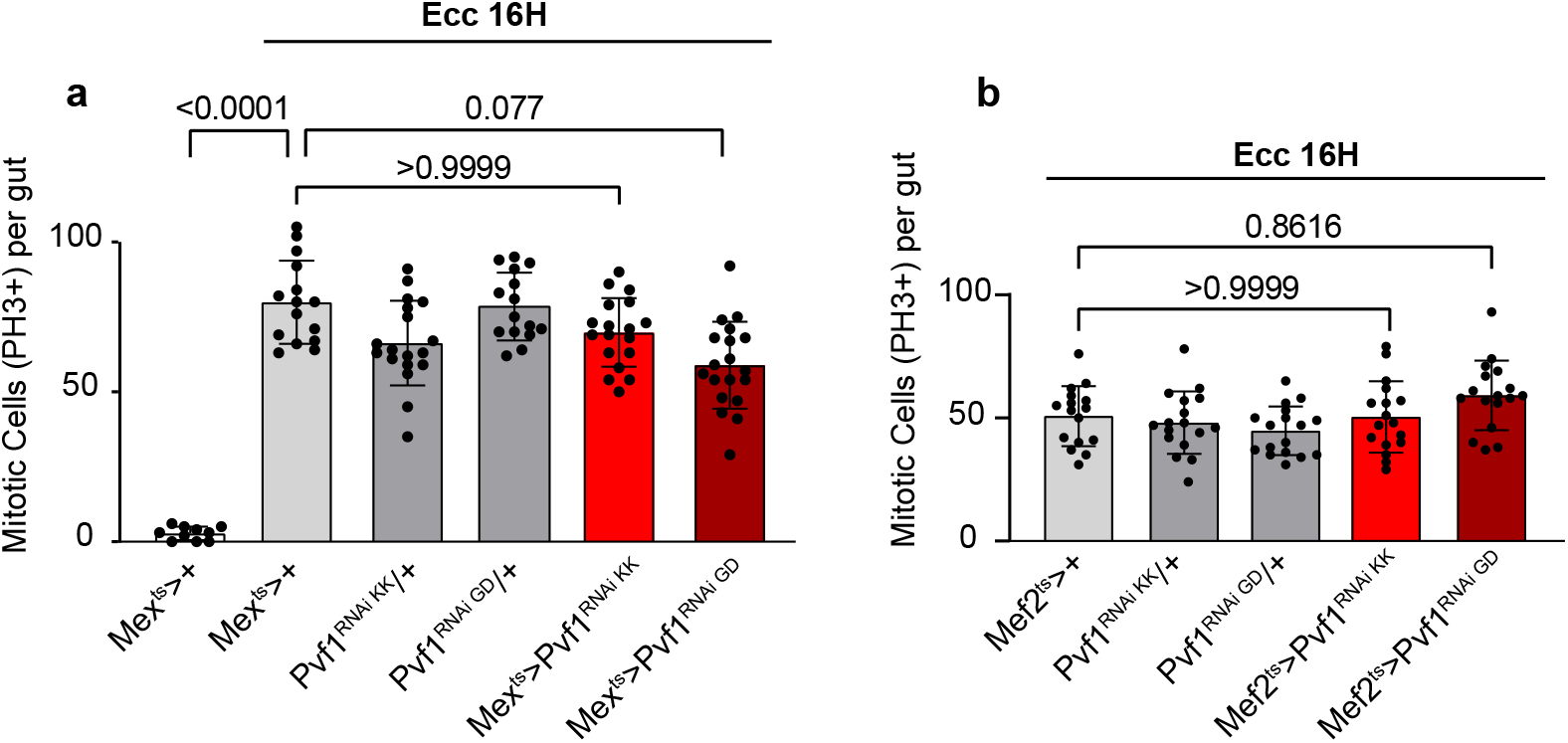
Pvf1 is not required in ECs or VMs for the infection-induced proliferative response. (a-b) Knock down of Pvf1 in ECs or VMs does not affect the infection-induced proliferative response. Quantification of PH3+ cells in midguts dissected from control flies or flies with EC-(a) or VM-specific (b) RNAi-mediated Pvf1 knockdown subjected to oral infection with Ecc15. Significance was tested with Kruskal Wallis with post-hoc multiple comparison analysis. Data are presented as mean values ±SD.

